## Supplementary Note 1 - Protocol Smart-seq2 for "Introducing synthetic thermostable RNase inhibitors to single-cell RNA-seq"

### Supplementary Note 1:
### Smart-seq2 with SEQURNA thermostable RNase inhibitor

In this version of the Smart-seq2 protocol, the use of a recombinant RNase inhibitor is replaced by the SEQURNA synthetic thermostable RNase inhibitor (RI). The overall protocol differs from the standard Smart-seq2 protocol only in that the SEQURNA RI is added only to the cell lysis buffer (i.e., cell collection buffer) and **not** supplemented again in the Reverse Transcription step. This is because the SEQURNA RI remains effective throughout the cell lysis and RNA denaturation step at 72°C and the following Reverse Transcription, while the protein-based Recombinant RNase inhibitor used in the original Smart-seq2 protocol, may denature and lose RI capacity during heating.

**Important notes:**

-Note that the optimal SEQURNA RI concentration in Smart-seq3 and is different than for Smart-seq2.

-Note that using more RNase inhibitor than indicated amounts is not beneficial. Excessive amounts of added inhibitor may result in decreased cDNA library yield and quality.

-Slight variations or modifications of the Smart-seq2 protocol are used in different labs, e.g., slight change in lysis buffer detergent or concentration: Use the suggested concentration of SEQURNA RI in the Smart-seq2 lysis buffer independent of Smartseq2 version.

-For a detailed protocol description and the subsequent sequencing library generation steps and indexing, refer to the original Smart-seq2 protocol:
Picelli 2014, Nature Protocols
https://www.nature.com/articles/nprot.2014.006

-For further details on the development of Smart-seq2, refer to the original Smartseq2 paper:
Picelli 2013, Nature Methods
https://www.nature.com/articles/nmeth.2639

-For in-depth information about the SEQURNA RNase inhibitor as well as potential updates of the protocol, please visit www.sequrna.com.

**Abbreviations**

DTT – Dithiothreitol
RI – RNase inhibitor
RT – Reverse Transcription
SS2 – Smart-seq2
TSO – Template-Switching Oligo

Oligonucleotide sequences (5’ to 3’):
SS2 oligo dT: 5′–AAGCAGTGGTATCAACGCAGAGTACT30VN-3′

SS2 TSO: 5′-AAGCAGTGGTATCAACGCAGAGTACATrGrG+G-3′
ISPCR: 5′-AAGCAGTGGTATCAACGCAGAGT-3′

### Prepare lysis plates

Prepare lysis buffer mix:
Note: Optimal concentration range of SEQURNA RI in the Smart-seq2 protocol is between 1-2 Mass U/µL in the lysis buffer, resulting in 0.45-0.9 Mass U/µL in the following RT step.

| Reagent | Conc. in lysis buffer | µL per reaction | 96 well plate (110 rxns) | 384 well plate (410 rxns) |
|---|---|---|---|---|
| 0.2% Triton X-100 | 0.08% | 1.9 | 209 | 779 |
| SEQURNA RI (50 Mass U/µL) | 1.2 Mass U/µL | 0.11 | 11 | 41 |
| dNTPs mix (10 mM) | 2.2 mM | 1 | 110 | 410 |
| SS2 oligo dT primer (10 µM) | 2.2 µM | 1 | 110 | 410 |
| Nuclease-free water | - | 0.49 | 53.9 | 200.9 |
| ERCC spike-ins (Optional) | - | - | - | - |
| Total | | 4.5 µL | 495 µL | 1845 µL |

Add 4.5 µL lysis buffer to each well of a 96/384 well plate, and centrifuge briefly to collect lysis buffer in the bottom of the wells.

### Sample collection

Sort single cells into 4.5 µL of lysis buffer lysis in either 96 or 384 wells.

Seal the plate with appropriate cover seals (tolerating -80°C to >100°C) and centrifuge the finished sorted plate immediately after. Transfer the plate to a -80°C freezer if not processing the cells into cDNA libraries within 1 day (keep plates in ~4°C fridge up to 1 day).

### Cell lysis

Remove the plate of sorted cells from the -80°C freezer and incubate in a thermocycler with heated lid at 72 °C for 3 min, followed by a 4 °C hold. Ensure that the plate is properly sealed, to avoid evaporation (use thermal pads, depending on thermocycler model).

### Reverse Transcription

While the plate is incubating at the cell lysis step, prepare the following Reverse transcription master-mix.

Note: Do **not** add additional inhibitor in the reverse transcription step. The SEQURNA RI from the lysis buffer stays effective throughout lysis and the following RT.

| Reagent | Reaction conc. | µL per reaction | 96 well plate (110 rxns) | 384 well plate (410 rxns) |
|---|---|---|---|---|
| SuperScript II reverse transcriptase (200 U/µL) | 100 U | 0.5 | 55 | 205 |
| Superscript II First Strand buffer (5x) | 1x | 2 | 220 | 820 |
| DTT (100 mM) | 5 mM | 0.5 | 55 | 205 |
| Betaine (5 M) | 1 M | 2 | 220 | 820 |
| $MgCl_2$ (1 M) | 10 mM | 0.1 | 11 | 41 |
| TSO (100 µM) | 1 µM | 0.1 | 11 | 41 |
| Nuclease-free water | | 0.3 | 33 | 123 |
| Total | | 5.5 µL | 605 µL | 2255 µL |

Add 5.5 µL RT mix to each well of a 96/384 well plate.

Replace the storage seal with a PCR seal. Ensure that the plate is properly sealed to avoid evaporation (use thermal pads, depending on thermocycler model).

Briefly centrifuge to collect reaction at the bottom of the tube.

Incubate the plate in a thermocycler at:

| Temp | Time | Cycles |
|---|---|---|
| 42 °C | 90 min | 1x |
| 50 °C | 2 min | 10x |
| 42 °C | 2 min | |
| 72 °C | 15 min | 1x |
| 4 °C | Hold | Hold |

### Preamplification PCR

Start preparing the PCR mix, when the incubation of the reverse transcription reaction is near completion, by combining the following components.

| Reagent | Reaction conc. | µL per reaction | 96 well plate (110 rxns) | 384 well plate (410 rxns) |
|---|---|---|---|---|
| First-strand reaction | – | 10 | 1100 | 4100 |
| KAPA HiFi HotStart ReadyMix (2x) | 1x | 12.5 | 1375 | 5125 |
| ISPCR primers (10 µM) | 0.08 µM | 0.2 | 22 | 82 |
| Nuclease-free water | – | 2.3 | 253 | 943 |
| Total volume | – | 25 µL | 2750 µL | 10250 µL |

Add 15 µL PCR mix to each well of a 96/384 well plate.

Briefly centrifuge to collect reaction at the bottom of the plate. Seal with a new PCR seal. Ensure that the plate is properly sealed, to avoid evaporation (use thermal pads, depending on thermocycler model).

Incubate the plate in a thermocycler at:

| Step | Temp | Time | Cycles |
|---|---|---|---|
| Initial denaturation | 98 °C | 3 min | 1x |
| Denaturation | 98 °C | 20 sec | 18-25x* |
| Annealing | 67°C | 15 sec | |
| Elongation | 72 °C | 6 min | |
| Final Elongation | 72 °C | 5 min | 1x |
| Hold | 4 °C | Hold | |

* depending on cell type (reflecting RNA content per cell)

### cDNA purification

Purification of cDNA is performed using Ampure XP beads or equivalent, e.g., 22% PEG Clean-up Beads (https://www.protocols.io/view/smart-seq3-protocol36wgq5rjxgk5 ).

1. To purify cDNA, add 0.8:1 ratio of beads to sample (20 µL) and mix by gently pipetting up and down.

2. Incubate at room temperature for 8 min.

3. Place on magnet and allow beads to settle ~5 min.

4. Discard supernatant, and wash pellets with freshly prepared 80% ethanol, keeping the plate on the magnet. The volume of ethanol will depend on the type of plate used (e.g. ~100 µL in case of 96-well plate).

5. Remove ethanol and repeat step 4.

6. Remove all ethanol and let the beads air dry for 2-5 min (do not over-dry the pellets).

7. Elute cDNA by adding 18 µL UltraPure Water or other suitable elution buffer (e.g., 10mM Tris-HCl, pH8.5) onto the pellets.

8. Remove the plate from the magnet and resuspend beads by pipetting up and down. Incubate for 8 min.

9. Place on magnet until clear (~3 min) and collect the eluate, containing the purified cDNA, to fresh plates or tubes.

Quality Control check

Inspect the cDNA library concentration and size distribution, e.g., on an Agilent Bioanalzyer High Sensitivity DNA Analysis chip.

Representative Bioanalyzer image of successfully amplified Smart-seq2 cDNA from a HEK cell using the SEQURNA RI:

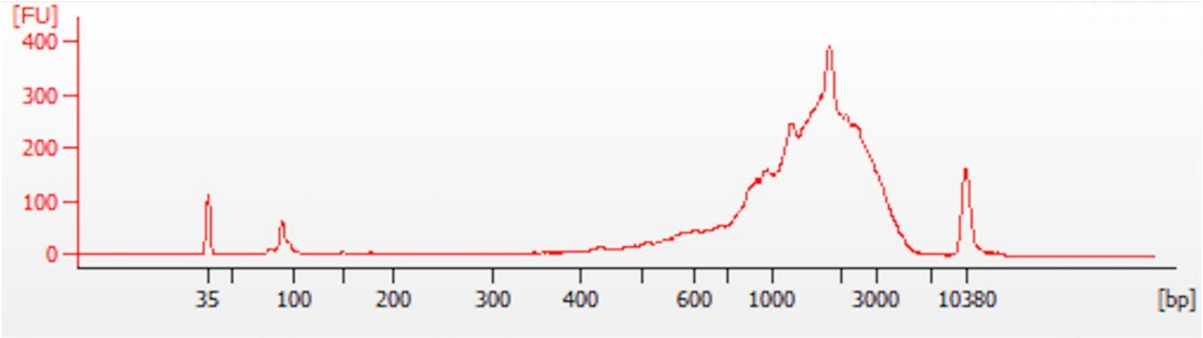


Trace of Smart-seq2 cDNA trace from a HEK cell, using an Agilent Bioanalzyer High Sensitivity DNA Analysis chip.

To prepare indexed sequencing libraries from Smart-seq2 cDNA by tagmentation and PCR, please refer to Picelli 2014, Nature Protocols: https://www.nature.com/articles/nprot.2014.006
