## Supplementary Note 2 - Protocol Smart-seq3 for "Introducing synthetic thermostable RNase inhibitors to single-cell RNA-seq"

### Supplementary Note 2
### Smart-seq3 with SEQURNA thermostable RNase inhibitor

In this version of the Smart-seq3 protocol, the use of a recombinant RNase inhibitor is replaced by the SEQURNA synthetic thermostable RNase inhibitor (RI). The overall protocol differs from the standard Smart-seq3 protocol only in that the SEQURNA RI is added only to the cell lysis buffer (i.e., cell collection buffer) and **not** supplemented again in the Reverse Transcription step. This is because the SEQURNA RI remains effective throughout the cell lysis and RNA denaturation step at 72°C and the following Reverse Transcription, while the protein-based Recombinant RNase inhibitor used in the original Smart-seq3 protocol, may denature and lose RI capacity during heating.

**Important notes:**

-Note that the optimal SEQURNA RI concentration in Smart-seq2 lysis buffer is different than for Smart-seq3.

-Note that using more RNase inhibitor than indicated amounts is not beneficial. Excessive amounts of added inhibitor may result in decreased cDNA library yield and quality.

-Slight variations or modifications of the Smart-seq3 protocol are used in different labs, e.g., slight change in lysis buffer detergent or concentration: Use the suggested concentration of SEQURNA RI in the Smart-seq3 lysis buffer independent of Smartseq3 version.

-For a detailed protocol description and the subsequent sequencing library generation steps and indexing, refer to the online protocol on protocols.io:
https://dx.doi.org/10.17504/protocols.io.bcq4ivyw

-For further details on the development of Smart-seq3, refer to the original Smartseq3 paper:
Hagemann-Jensen 2020, Nature Biotechnology
https://www.nature.com/articles/s41587-020-0497-0

-For in-depth information about the SEQURNA RNase inhibitor as well as potential updates of the protocol, please visit www.sequrna.com.

**Abbreviations**

DTT – Dithiothreitol
RI – RNase inhibitor
RT – Reverse Transcription
SS3 – Smart-seq3
TSO – Template-Switching Oligo

Oligonucleotide sequences (5’ to 3’):
SS3 oligo dT: 5′-/5Biosg/ACGAGCATCAGCAGCATACGAT30VN-3′

SS3 TSO: 5′-/5Biosg/AGAGACAGATTGCGCAATGNNNNNNNNrGrGrG-3′
SS3 Fwd Primer:
5′-TCGTCGGCAGCGTCAGATGTGTATAAGAGACAGATTGCGCAA*T*G-3′
SS3 Rev Primer: 5′-ACGAGCATCAGCAGCATAC*G*A-3′
* phosphorothioate bonds

### Prepare lysis plates

Prepare lysis buffer mix:
Note: Optimal concentration of SEQURNA RI in the Smart-seq3 protocol is 0.15-0.3 Mass U/µL in the lysis buffer, resulting in 0.11-0.23 Mass U/µL in the following RT step.

| Reagent | Conc. in lysis buffer | µL per reaction | 96 well plate (110 rxns) | 384 well plate (410 rxns) |
|---|---|---|---|---|
| Poly-ethylene Glycol 8000 (50% solution) | 6.7% | 0.40 | 44 | 164 |
| Triton X-100 (10% solution) | 0.1% | 0.03 | 3.3 | 12.3 |
| SEQURNA Inhibitor (50 Mass U/µL) | 0.2 Mass U/µL | 0.012 | 1.32 | 4.92 |
| SS3 oligo dT (100µM) | 0.67µM | 0.02 | 2.2 | 8.2 |
| dNTPs (25mM/each) | 0.67 mM/each | 0.08 | 8.8 | 32.8 |
| Nuclease Free Water | - | 2.46 | 270.6 | 1008.6 |
| ERCC spike-ins (Optional) | - | - | - | - |
| Total | - | 3 µL | 330 µL | 1230 µL |

Add 3 µL lysis buffer to each well of a 96/384 well plate, and centrifuge briefly to collect lysis buffer in the bottom of the wells.

### Sample collection

Sort single cells into 3 µL of lysis buffer lysis in either 96 or 384 wells.

Seal with appropriate seals (tolerating -80°C to >100°C) and centrifuge the finished sorted plate immediately after. Transfer the plate to a -80°C freezer if not processing the cells into cDNA libraries within 1 day (keep plates in ~4°C fridge up to 1 day).

### Cell lysis

Remove the plate of sorted cells from the -80°C freezer and incubate in a thermocycler with heated lid at 72 °C for 3 min, followed by a 4 °C hold. Ensure that the plate is properly sealed, to avoid evaporation (use thermal pads, depending on thermocycler model).

### Reverse Transcription

While the plate is incubating at the cell lysis step, prepare the following Reverse transcription master-mix.

Note: Do **not** add additional inhibitor in the reverse transcription step. The SEQURNA RI from the lysis buffer stays effective throughout lysis and the following RT.

| Reagent | Conc. in RT | µL per reaction | 96 well plate (110 rxns) | 384 well plate (410 rxns) |
|---|---|---|---|---|
| Tris-HCl pH 8.3 (1M) | 25mM | 0.1 | 11 | 41 |
| NaCl (1M) | 30mM | 0.12 | 13.2 | 49.2 |
| $MgCl_2$ (100mM) | 2.5mM | 0.1 | 11 | 41 |
| GTP (100mM) | 1mM | 0.04 | 4.4 | 16.4 |
| DTT (100mM) | 8mM | 0.32 | 35.2 | 131.2 |
| SS3 TSO (100µM) | 2µM | 0.08 | 8.8 | 32.8 |
| Maxima H-minus RT enzyme (200U/µL) | 8U | 0.04 | 4.4 | 16.4 |
| Nuclease Free Water | - | 0.2 | 22 | 82 |
| Total | | | | |
| | - | 1µL | 110µL | 410µL |

Add 1 µL RT mix to each well of a 96/384 well plate.

Replace the storage seal with a PCR seal. Ensure that the plate is properly sealed to avoid evaporation (use thermal pads, depending on thermocycler model).

Briefly centrifuge to collect reaction at the bottom.

Incubate the plate in a thermocycler at:

| Temp | Time | Cycles |
|---|---|---|
| 42 °C | 90 min | 1x |
| 50 °C | 2 min | 10x |
| 42 °C | 2 min | |
| 85 °C | 5 min | 1x |
| 4 °C | Hold | Hold |

### Preamplification PCR

Start preparing the PCR mix, when the incubation of the reverse transcription reaction is near completion, by combining the following components.

| Reagent | Conc. in PCR | µL per reaction | 96 well plate (110 rxns) | 384 well plate (410 rxns) |
|---|---|---|---|---|
| Kapa HiFi HotStart buffer (5X) | 1X | 2.0 | 220 | 820 |
| dNTPs (25mM/each) | 0.3mM/each | 0.12 | 13.2 | 49.2 |
| $MgCl_2$ (100mM) | 0.5mM | 0.05 | 5.5 | 20.5 |
| Fwd Primer (100µM) | 0.5µM | 0.05 | 5.5 | 20.5 |
| Rev Primer (100µM) | 0.1µM | 0.01 | 1.1 | 4.1 |
| Kapa Polymerase (1U/µL) | 0.02U/µL | 0.2 | 22 | 82 |
| Nuclease Free Water | – | 3.57 | 392.7 | 1463.7 |
| Total | – | 6µL | 660µL | 2460µL |

Add 6 µL PCR mix to each well of a 96/384 well plate.

Briefly centrifuge to collect reaction at the bottom. Seal with a new PCR seal. Ensure that the plate is properly sealed to avoid evaporation (use thermal pads, depending on thermocycler model).

Incubate the plate in a thermocycler at:

| Step | Temp | Time | Cycles |
|---|---|---|---|
| Initial denaturation | 98 °C | 3 min | 1x |
| Denaturation | 98 °C | 20 sec | 18-25x* |
| Annealing | 65°C | 30 sec | |
| Elongation | 72 °C | 4 min | |
| Final Elongation | 72 °C | 5 min | 1x |
| Hold | 4 °C | Hold | |

* depending on cell type (reflecting RNA content per cell)

### cDNA purification

Purification of cDNA is performed using Ampure XP beads or equivalent, e.g., 22% PEG Clean-up Beads (https://www.protocols.io/view/smart-seq3-protocol36wgq5rjxgk5 ).

1. To purify cDNA add 0.8:1 ratio of 22% PEG beads to sample, and mix by gently pipetting up and down.

2. Incubate at room temperature for 8 min.

3. Place on magnet and allow beads to settle for ~5 min.

4. Discard supernatant, and wash once with 20 µL/ 100 µL of freshly prepared 80% ethanol for 384 / 96 well plates respectively, keeping the plate on the magnet.

5. Remove ethanol and let the beads air dry for 2-5 min (do not over-dry the pellets).

6. Elute cDNA in 12 µL of UltraPure Water or other suitable elution buffer (e.g., 10mM Tris-HCl, pH8.5) onto the pellets.

7. Remove the plate from the magnet and resuspend beads by pipetting up and down. Incubate for 8 min.

8. Place on magnet until clear (~3 min) and collect the eluate, containing the purified cDNA, to fresh plates or tubes.

Quality Control check

Inspect the cDNA library concentration and size distribution, e.g., on an Agilent Bioanalzyer High Sensitivity DNA Analysis chip.

Representative Bioanalyzer image of successfully amplified Smart-seq3 cDNA from a HEK cell using the SEQURNA RI:

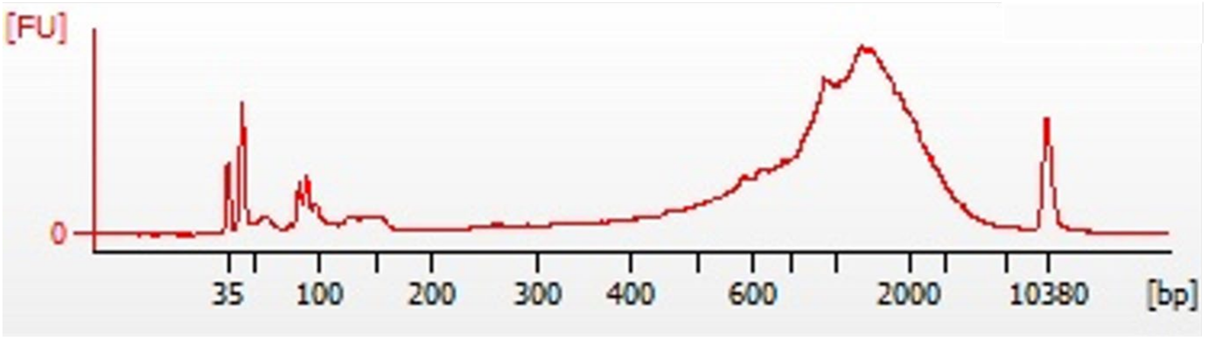


Trace of Smart-seq3 cDNAfrom a HEK cell, using an Agilent Bioanalzyer High Sensitivity DNA Analysis chip.

To prepare indexed sequencing libraries from Smart-seq3 cDNA by tagmentation and PCR, please refer to online protocol https://dx.doi.org/10.17504/protocols.io.bcq4ivyw.
