## Supplementary Note 3 - Protocol Smart-seq3xpress for "Introducing synthetic thermostable RNase inhibitors to single-cell RNA-seq"

### Supplementary Note 3:
### Smart-seq3xpress with SEQURNA thermostable RNase inhibitor

In this version of the Smart-seq3xpress protocol, the use of a recombinant RNase inhibitor is replaced by the SEQURNA synthetic thermostable RNase inhibitor (RI). The overall protocol differs from the standard Smart-seq3xpress protocol only in that the SEQURNA RI is added only to the cell lysis buffer (i.e., cell collection buffer) and **not** supplemented again in the Reverse Transcription step. This is because the SEQURNA RI remains effective throughout the cell lysis and RNA denaturation step at 72°C and the following Reverse Transcription, while the protein-based Recombinant RNase inhibitor used in the original Smart-seq3xpress protocol, may denature and lose RI capacity during heating.

**Important notes:**
-Note that using more RNase inhibitor than indicated amounts is not beneficial. Excessive amounts of added inhibitor may result in decreased library yield and quality.

-This protocol requires a liquid handler capable of nanoliter dispenses.

- For a detailed protocol description and the subsequent sequencing library generation steps, refer to the online protocol on protocols.io:
https://www.protocols.io/view/smart-seq3xpress-yxmvmk1yng3p/v2

-For further details on the development of Smart-seq3xpress, refer to the original Smart-seq3xpress paper:
Hagemann-Jensen 2022, Nature Biotechnology
https://www.nature.com/articles/s41587-022-01311-4

-For in-depth information about the SEQURNA RNase inhibitor, please refer to our white paper available at www.sequrna.com.

**Abbreviations**
DTT – Dithiothreitol
RI – RNase inhibitor
RT – Reverse Transcription
TSO – Template-Switching Oligo

Oligonucleotide sequences (5' to 3'):
SS3 oligo dT: 5′-/5Biosg/ACGAGCATCAGCAGCATACGAT30VN-3′
Smart-seq3xpress TSO: 5′-
/5BiosG/AGAGACAGATTGCGCAATGNNNNNNNNWWrGrGrG-3′
SS3 Fwd Primer: 5′-
TCGTCGGCAGCGTCAGATGTGTATAAGAGACAGATTGCGCAA*T*G-3′
SS3 Rev Primer: 5′-ACGAGCATCAGCAGCATAC*G*A-3′
* phosphorothioate bonds

### Prepare overlay plates

Use Vapor-Lock, Sillicone oil 25 cSt, Silicone Oil 100 cSt (The higher viscosity is better suited for shipping plates). CAUTION: Do not dispense these silicone oils / overlays with your non contact liquid handler. The solutions can "creep" everywhere. Use either manual multichannel pipettes or semi-manual (e.g. Integra ViaFlow) / automatic dispensing (e.g. Agilent Bravo, Tecan Fluent) with tips, and prepare and store in bulk.

Add 3 µL of overlay to each well of a 384 well plate. The amount of overlay can be increased if desired.

Quick pulse centrifugation to 1000 x g to ensure all is collected in the bottom of wells.

Put on seal and store at Room temperature until use.

### Prepare lysis plates

Prepare lysis buffer mix:
Note: Optimal concentration of SEQURNA RI in the Smart-seq3xpress lysis buffer is 0.2 Mass U/µL, resulting in 1.5 Mass U/µL in the following RT step.

| Reagent | Conc. in lysis buffer | µL per reaction | 384 well plate (500 rxns) |
|---|---|---|---|
| Poly-ethylene Glycol 8000 (40% solution) | 6.7% | 0.05 | 25 |
| Triton X-100 (10% solution) | 0.1% | 0.003 | 1.5 |
| SEQURNA Inhibitor (50 Mass U/µL) | 0.2 Mass U/µL | 0.0012 | 0.6 |
| SS3 oligo dT (10µM) | 0.167µM | 0.005 | 2.5 |
| dNTPs (10mM/each) | 0.66mM/each | 0.02 | 10 |
| Nuclease Free Water | - | 0.221 | 110.5 |
| ERCC spike-ins (Optional) | - | - | - |
| Total | - | 0.3 µL | 150 µL |

Add 0.3 µL lysis buffer to each well of a 384 well plate containing overlay, and centrifuge briefly to collect lysis buffer.

### Sample collection

Sort single cells into 0.3 µL of lysis buffer with overlay in 384 wells.

Seal with appropriate seals (tolerating -80°C to >100°C) and centrifuge the finished sorted plate immediately. Transfer the plate to a -80°C freezer if not processing the cells into cDNA libraries within 1 day (or keep plates in ~4°C fridge up to 1 day).

### Cell lysis

Remove the plate of sorted cells from the -80°C freezer and incubate in a thermocycler with heated lid at 72 °C for 10 min, followed by a 4 °C hold.

### Reverse Transcription

While the plate is incubating at the cell lysis step, prepare the following Reverse transcription master-mix.

*Note:* Do **not** add additional inhibitor in the reverse transcription step. The SEQURNA RI from the lysis buffer stays effective throughout lysis and the following RT.

| Reagent | Conc. in RT | µL per reaction | 384 well plate (500 rxns) |
|---|---|---|---|
| Tris-HCl pH 8.3 (1M) | 25mM | 0.01 | 5 |
| NaCl (2.5M) | 30mM | 0.0048 | 2.4 |
| MgCl2 (100mM) | 2.5mM | 0.01 | 5 |
| GTP (100mM) | 1mM | 0.004 | 2 |
| DTT (100mM) | 8mM | 0.032 | 16 |
| Smart-seq3xpress TSO (100µM) | 0.75µM | 0.003 | 1.5 |
| Maxima H-minus RT enzyme (200U/µL) | 2U | 0.004 | 2 |
| Nuclease Free Water | - | 0.0325 | 16.25 |
| Total | - | 0.1µL | 50µL |

Add 0.1 µL RT mix to each well of a 384 well plate.

Replace the storage seal with a PCR seal. Ensure that the plate is properly sealed, to avoid evaporation.

Briefly centrifuge to collect reaction at the bottom.

Incubate the plate in a thermocycler at:

| Temp | Time | Cycles |
|---|---|---|
| 42 °C | 90 min | 1x |
| 50 °C | 2 min | 10x |
| 42 °C | 2 min | |
| 85 °C | 5 min | 1x |
| 4 °C | Hold | Hold |

### Preamplification PCR

Start preparing the PCR mix, when the incubation of the reverse transcription reaction is near completion, by combining the following components.

| Reagent | Reaction conc. | µL per reaction | 384 well plate (500 rxns) |
|---|---|---|---|
| SeqAmp PCR buffer (2x) | 1X | 0.5 | 250 |
| Fwd Primer (100µM) | 0.5µM | 0.005 | 2.5 |
| Rev Primer (100µM) | 0.5µM | 0.005 | 2.5 |
| SeqAmp DNA polymerase (1.25u/uL) | 0.025U/µL | 0.02 | 10 |
| Nuclease Free Water | – | 0.07 | 35 |
| Total | – | 0.6µL | 300µL |

Add 0.6 µL PCR mix to each well of a 384 well plate.

Briefly centrifuge to collect reaction at the bottom.

Incubate the plate in a thermocycler at:

| Step | Temp | Time | Cycles |
|---|---|---|---|
| Initial denaturation | 98 °C | 1 min | 1x |
| Denaturation | 98 °C | 10 sec | 12-16x* |
| Annealing | 65°C | 30 sec | |
| Elongation | 72 °C | 4 min | |
| Final Elongation | 72 °C | 10 min | 1x |
| Hold | 4 °C | Hold | |

* depending on cell type (reflecting RNA content per cell)

To prepare indexed sequencing libraries from Smart-seq3express cDNA by tagmentation and PCR, please refer to online protocol: https://www.protocols.io/view/smart-seq3xpress-yxmvmk1yng3p/v2.

#### Quality Control check

Inspect final library concentration and size distribution after tagmentation, PCR, and clean-up, e.g., on an Agilent Bioanalzyer High Sensitivity DNA Analysis chip.

Representative Bioanalyzer image of a successfully tagmented and pooled library using the SEQURNA RNase inhibitor:

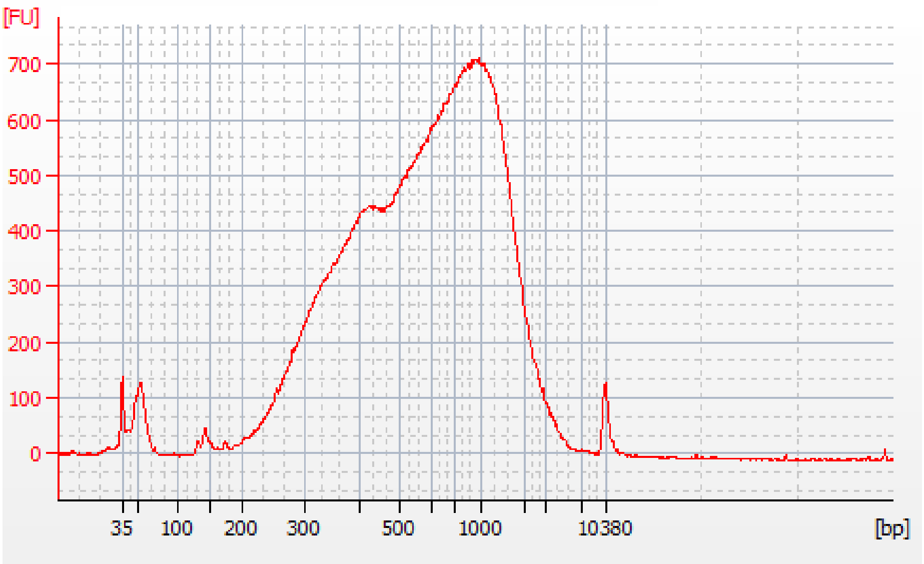

Trace of Smart-seq3express pooled library (after tagmentation and PCR amplification) from sorted HEK cells, using an Agilent Bioanalzyer High Sensitivity DNA Analysis chip.
